# Organ-specific prioritization and annotation of non-coding regulatory variants in the human genome

**DOI:** 10.1101/2023.09.07.556700

**Authors:** Nanxiang Zhao, Rintsen N. Sherpa, Shengcheng Dong, Kobe B. Howcroft, Alan P. Boyle

## Abstract

Identifying non-coding regulatory variants in the human genome remains a challenging task in genomics. Recently, we released the second version of our leading regulatory variant database, RegulomeDB. Building upon this comprehensive database, we developed a novel machine-learning architecture, TLand, which utilizes RegulomeDB-derived features to predict regulatory variants at the cell- or organ-specific level. In our holdout benchmarking, TLand consistently outperformed state-of-the-art models, demonstrating its ability to generalize to new cell lines or organs. We trained three types of organ-specific TLand models to overcome the common model bias toward high data availability cell lines or organs. These models accurately prioritize relevant organs for 2 million GWAS SNPs associated with GWAS traits. Moreover, our analysis of top-scoring variants in specific organ models showed a high enrichment of relevant GWAS traits. We expect that TLand and RegulomeDB will further advance our ability to understand human regulatory variants genome-wide.

## Background

Understanding the biological impact of variants located in non-coding regions of the human genome is a significant challenge. Nearly 90% of disease-risk-associated single-nucleotide polymorphisms (SNPs) identified from genome-wide association studies (GWAS) are within non-coding regions. Similarly, 75% of patients with Mendelian disease have mutations outside of protein-coding regions (1). Delving into the functional consequences of these disease-associated non-coding variants may uncover potential causal SNPs, expanding our insights beyond those within the coding region.

Prioritizing non-coding variants requires integrating multiple layers of functional information, including regulatory annotations identified from high-throughput sequencing datasets (e.g., DNase-seq (2), ChIP-seq (3), and ATAC-seq (4)). Such annotations provide additional information for pinpointing causal variants, which are often not the lead variants identified in GWAS. Despite the benefit of incorporating functional genomics assay-based evidence when examining non-coding variants, the lack of available variant annotation tools limits the use of such data. Most resources developed for clinical purposes have focused on coding regions as an application of exome sequencing-based data (5,6), which captures less than 5% of human variation (7–9).

Previously, we developed RegulomeDB, a comprehensive database for prioritizing and annotating variants in non-coding regions, which has been widely used in the research community (10). RegulomeDB intersects query variants with regulatory regions predicted by functional genomics assays and, by utilizing ranking heuristics, informs users about putative functional consequences to prioritize variants. Recently, RegulomeDB has been upgraded to version 2 (11), improving its annotation power by incorporating thousands of new functional genomics datasets from the ENCODE project (12), Roadmap Epigenomics Consortium (13), and the Genomics of Gene Regulation Consortium. A suite of models, namely SURF and TURF, was developed and integrated in this version to provide accurate probabilistic scores for general and cell-type specific regulatory activities (14,15).

However, a drawback of this model suite is that it was trained using hg19-referenced ENCODE datasets for only six common cell lines. The resulting models are biased and lack sufficient statistical power to generalize their predictions to less-studied cell lines. As a result, RegulomeDB scores are less informative for cell lines, tissues, and organs that lack the abundance of data available for commonly-studied cell lines. This can make it challenging for users to identify variants of interest and formulate hypotheses regarding their regulatory functions. Nonetheless, we anticipate that RegulomeDB will continue to improve in the years to come as new datasets are released by the Encyclopedia of DNA Elements (ENCODE) (12) and Impact of Genomic Variation on Function (IGVF) (16) consortia, spanning additional cell lines, tissues, and assay types, such as Hi-C (17) and Enformer predictions (18). However, the current SURF and TURF models were not designed to incorporate data continuously, and are potentially prone to overfitting as the feature space expands due to the simple model architecture. Given the breadth and volume of functional genomics datasets being released each day, these are serious limitations.

Here, we present TLand, a flexible architecture based on stacked generalization (19) that learns RegulomeDB-derived features to predict regulatory variants at the cell- or organ-specific level. TLand’s stacked generalization approach groups features into biologically meaningful subspaces, training individual estimators before assembly to reduce overfitting and enable further integration of features. Cell-specific TLand consistently outperformed state-of-the-art models in benchmarking with a hold-out cell line. By developing a suite of models rather than a single, monolithic model, organ-specific TLand addresses the data availability bias and further improves on cell-specific TLand in identifying regulatory variants relevant to the target organ, as predicted by GWAS. Furthermore, analysis of the top-scoring variants in specific organ models revealed a high enrichment for correlated GWAS traits. Given its superior performance relative to its predecessors and competing methods, we expect TLand to address the ongoing challenge of reliably prioritizing variants, even in less-studied cell lines and organs, thus advancing our ability to identify regulatory variants genome-wide.

## Methods

### Allele-specific binding (ASB) variants

We define a variant as likely-regulatory if it shows evidence for allele-specific binding (ASB), the molecular signature of which is significantly different when ChIP-seq reads are phased between the two alleles in a heterozygous individual. We trained our models on ASB variants for all TFs with ChIP-seq data in ENCODE. We included a total of 7,530 ASB variants in 6 cell lines (GM12878, HepG2, A549, K562, MCF7, and H1hESC) called by *AlleleDB* (20). We utilized negative training sets previously produced in our lab (15), encompassing non-ASB variants and randomly-selected background variants. In total, we included 14,773 unique variants in our training set. The complementary ASB data of IMR-90 and H9 used for evaluation were downloaded from Adastra (21) at https://adastra.autosome.org/bill-cipher/search/advanced?fdr=0.05&es=0&cl=IMR90%20(lung%20fibroblasts) and https://adastra.autosome.org/bill-cipher/search/advanced?fdr=0.05&es=0&cl=H9.

### Model architecture

TLand is a one-layer stacked architecture that consists of two parts: three base classifiers and one meta-classifier (Fig 1b and Supplementary Fig 1). TLand takes input as 19 generic features, 40 deep learning prediction-derived features, and 5 cell-specific features or 13 organ-specific features (Supplementary Table 1), which are directly derived from RegulomeDB queries. Zeros would be assigned if no hits were found in RegulomeDB queries. Features were bagged into 3 subspaces: experimental set (generic features and cell/organ-specific features), deep learning set (deep learning features and cell/organ-specific features), and cell/organ-specific set (only cell/organ-specific features). We selected lightGBM (22), random forest (23), and neural network (24) as base classifiers due to their distinct decision boundaries. Our base models were fine-tuned with Optuna (25). We used 300 estimators in random forest models, 250 boost rounds, and a 0.049 learning rate in lightGBM models, 3 layers with 128 neurons per layer, batch size of 128, adaptive learning rate, and with the max iteration of 30 in neuron network models. Probabilities were used as the output of base classifiers and to train our meta-classifier. We calculated interaction terms of probabilities up to the degree of 2 before feeding into our meta-classifier. We customized a ridge classifier to output probabilities with hyperparameter alpha as 1.9 as our final meta-classifier. The ridge classifier was trained with 4-fold group cross-validation. We grouped the training data based on the genomic positions (i.e., variants at the same genomic positions regardless of whether cell lines were in the same group). All models were specified with balanced class weights. LightGBM was implemented with the Python package (22). Random forest, neural network, and relevant pipelines were implemented with scikit-learn (26). The stacked generalization algorithm was implemented with mlxtend (27).

**Figure 1.**
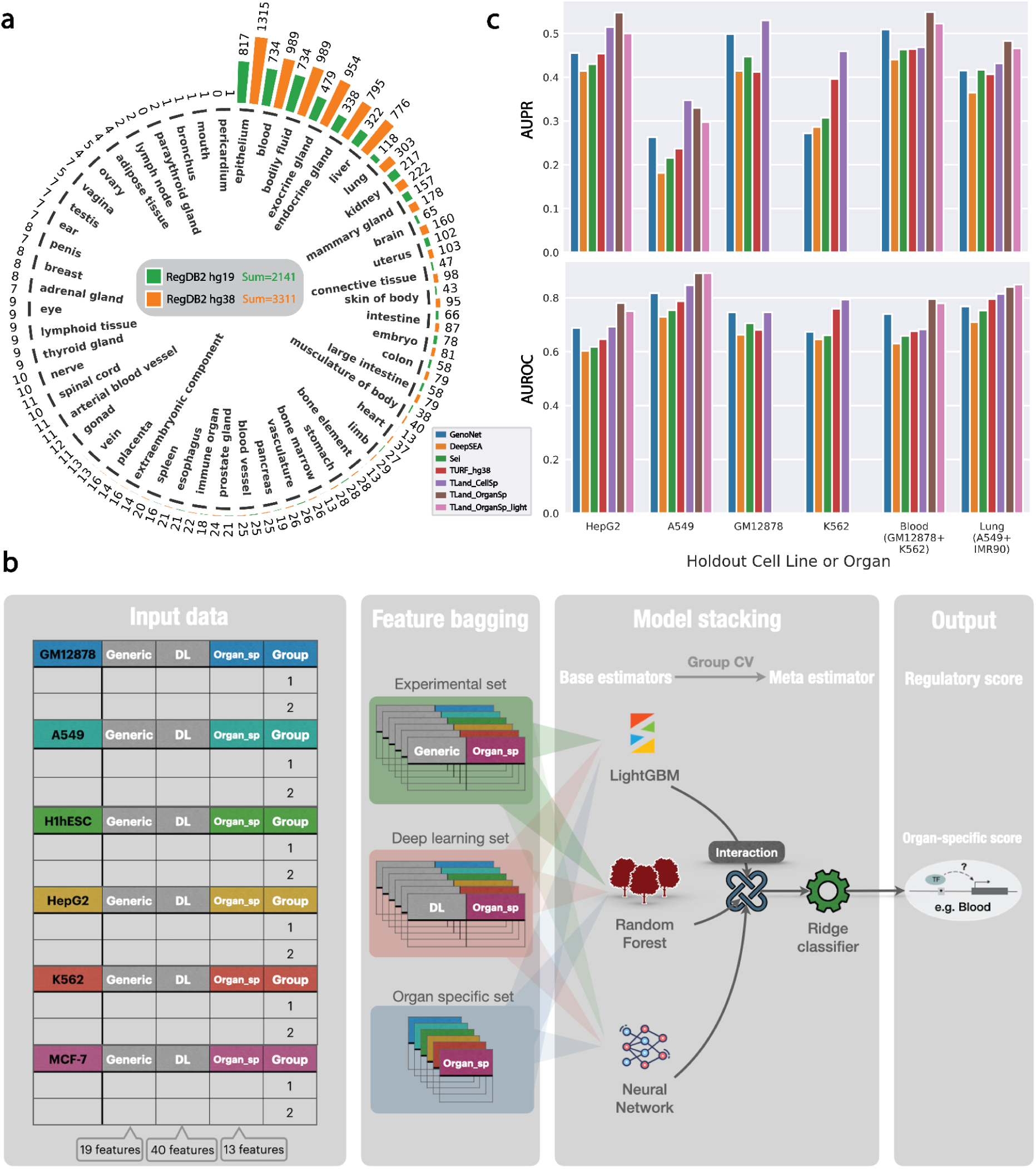
TLand improves regulatory variant predictions. (a) TF ChIP-seq data availability across organs on RegulomeDB v2. Green bar plots represent counts on GRCh38. Orange bar plots represent counts on hg19. The total number of counts for each assembly is in the middle of the gray box. Notice that the summation is not simply adding all numbers together due to some cell lines having multiple corresponding organs. (b) Organ-specific TLand architecture. Organ-specific TLand was trained to predict human regulatory variants in an organ-specific manner by using RegulomeDB-derived features. (c) Benchmarking TLand performance by AUROC and AUPR. The X-axis is holdout cell lines or organs. The Y-axis is AUPR on the top panel and AUROC on the bottom panel.

### Model training and evaluation

TLand was trained, validated, and tested on the generated ASB datasets. After concatenating across six cell lines, 88,638 (i.e., 14,773 x 6) variants were used for training and validation. The concatenation allowed us to have more data per prediction task. For the task of predicting an unseen cell line, we held out data for one cell line as a test case and then used the rest as training data for the TLand model. Simultaneously, during the train and test split, we ensured that unique variants, irrespective of cell lines, were exclusively present either in the training or test dataset. This precaution aimed to prevent information leakage, as features across cell lines are often correlated. The test and train split ratio for negative data is ⅙. Similarly, for organ-specific models, we held out one organ and unique variants as a test dataset and used organ-specific labels for splitting the data. We calculated both the Area Under the Receiver Operating Characteristic Curve (AUROC) and the Area Under the Precision Recall Curve (AUPR) to evaluate the classifiers. We also performed a separate evaluation using leave-one-chromosome-out cross-validation to account for any sequence-specific data leakage, as the Sei-derived features are conditional on 4096bp of sequence context. We aggregated the variant sets across cell lines, calculated the AUPR for each held out chromosome set, reported mean AUPR, and computed 95% confidence intervals using 1000 bootstrapped samples. After evaluations, the final models were trained with all 88,638 variants.

### DNase signal quantile normalization

All 1372 DNase bigwig default files processed on GRCh38 assembly (up to May 2022) were obtained from ENCODE (Supplementary Table 5). We designed an efficient pipeline to quantile normalize all signals to achieve the balance between accuracy, runtime, and storage (Supplementary Fig 2). BigWig files were first converted to BedGraph format using bigWigToBedGraph (28). Average signals were extracted for 10bp, non-overlapping windows extracted from each BedGraph file with bedmap (29) and stored in bed files. These bed files were concatenated and converted to parquet format for processing with qnorm (30), a Python package that applies a multithreaded incremental normalization strategy to efficiently normalize extremely large datasets (30). We quantile normalized 10-bp average signals for all 1372 DNase datasets with a batch size of 343 files per normalization iteration. Details of storage, runtime, and memory are shown in Supplementary Fig 2.

### Benchmarking other models and organ definition

We benchmarked TLand with state-of-the-art models, including GenoNet, DeepSEA, Sei, and TURF. Pre-calculated GenoNet scores were downloaded from https://zenodo.org/record/3336209/files/. We imputed the missing prediction scores of GenoNet as the average of all available scores. DeepSEA and Sei models were downloaded from the original paper (31,32). We ran the predictions locally. The previous best model, TURF, was retrained to predict regulatory variants on GRCh38 by querying and inputting the GRCh38 features (15). TURF predictions for unseen cell lines were derived by training separate models on each of the seen (i.e., training) cell lines, then taking the average of all trained models as an estimate for the unseen cell line. This resembles the ensemble method of majority vote. For organ-level predictions, we averaged relevant cell line model predictions to estimate the predictions, using the human organ definitions from ENCODE (https://www.encodeproject.org/summary/?type=Experiment&control_type!=*&replicates.library.biosample.donor.organism.scientific_name=Homo+sapiens&status=released.). For example, we averaged TURF model predictions of A549 and IMR-90 to predict lung regulatory variants.

### Evaluation with MPRA data

We download the new MPRA data of five cell lines from ENCODE, including GM12878 (ENCODE Accession ID: ENCSR560QIE), K562 (ENCSR971PLA), A549 (ENCSR793OLW), HCT116 (ENCSR873QHK), and SK-N-SH (ENCSR579QGV). We trained the final set of TLand models using all available data (i.e., from six training cell lines). Then we evaluated the final models by calculating the AUROC and AUPR given the cell line’s p-value threshold (Skew -log10(FDR) at least > 5) to define positive and negative labels.

### GWAS catalog variants and LD data

We first downloaded all known SNV positions significantly associated with traits described in the GWAS Catalog (https://www.ebi.ac.uk/gwas/api/search/downloads/alternative) (33). We then completed linkage disequilibrium (LD) expansion for each SNV, incorporating all SNVs from the 1000 Genome Project in strong LD (R^2^ threshold of 0.6). R^2^ values were downloaded from (gs://genomics-public-data/linkage-disequilibrium). In total, we calculated their TLand model scores for 1,974,549 SNVs.

Summary statistics for 94 GWAS traits from the UK BioBank (34) were downloaded from https://console.cloud.google.com/storage/browser/finucane-requester-pays/ukbb-finemapping. Metadata for these traits (trait definitions and number of samples, cases, and controls) were downloaded from https://www.finucanelab.org/data. Summary statistics were prepared for use in LDSC using the munge_sumstats tool from https://github.com/belowlab/ldsc/tree/2-to-3, and LD scores were calculated for each organ-specific TLand score and trait combination using the ldsc tool from https://github.com/bulik/ldsc.

### Target organs annotations for GWAS traits

Gold-standard target-organ associations for 44 GWAS traits were annotated with Open Targets (https://platform.opentargets.org/), EMBL-EBI Ontology Lookup Service (https://www.ebi.ac.uk/ols/index), and by reviewing literature for selected GWAS traits (Supplementary Table 3). These annotations were used to evaluate model performance for each GWAS trait.

### Prioritization of relevant organs for GWAS traits

We predicted approximately 2 million GWAS variants and strongly linked SNPs across 51 organs. For each manually annotated GWAS trait, we selected all associated SNPs. For each organ, we calculated the p-value from a one-tailed Student’s t-test (35) comparing the sampled organ-specific SNP scores versus the population distribution, defined as the 2 million SNP score distribution for the organ. We ranked organs by the inverse of their p-values, using a significance threshold of 0.05 to select only organs with statistical support for a functional association. We evaluated performance by comparing the ranked (i.e., prioritized) organs with the manually annotated target organs for each trait. Accuracy was defined as the fraction of correct overlaps between model-based and gold-standard organ annotations among the top 4 organs for each model. We combined any two models’ ranking lists by taking the union of their lists then ranked by p-value, and selected the top 5 organs for each combination.

### GWAS trait enrichment

We began by gathering the top-scoring GWAS variants, specifically those with organ-specific scores exceeding 0.5 for each organ model. We traced back from these variants to identify their associated traits. For each trait, we defined and computed the trait score as the sum of the organ-specific scores of the associated variants. To refine our results for each organ, we excluded traits with low total counts, where total counts represent the sum of trait counts for all associated GWAS and strongly-linked SNPs. Finally, we calculated organ-specific GWAS trait enrichment scores by normalizing the trait score to the total counts.

### Stratified LD Score Regression (S-LDSC)

S-LDSC (36) partitions complex trait heritability by genomic and functional annotations and is used to help identify which annotations are most relevant to the genetic architecture of a given trait. It estimates per-SNP heritability by regressing GWAS test statistics on LD scores that are stratified by these annotations.

The expected GWAS χ^2^ statistic of SNP *j* is modeled as

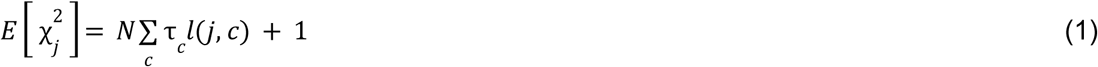

where *N* is the sample size, *C* is the set of annotations indexed by *c* (referred to as categories in the S-LDSC paper), *τ_c_* is the per-SNP contribution to heritability by annotation *C_c_*, and *l*(*j*,*c*) is the LD score of SNP *j* with respect to annotation *C_c_*. Enrichment of an annotation for a given trait is calculated using

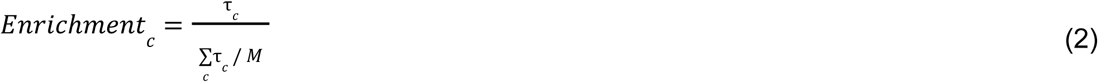

where *M* is the total number of SNPs used in the computation. This is interpreted as the proportion of SNP heritability contributed by an annotation divided by the proportion of SNPs in that annotation.

TLand scores for the 51 organs were calculated for all HapMap 3 SNPs (37) excluding those within the major histocompatibility complex (MHC) region of the genome (1,217,311 SNPs). TLand scores were binarized at a threshold of 0.7, covering an average of 1% of SNPs across organs. We performed S-LDSC (https://github.com/bulik/ldsc) for each of the 94 UKBB traits conditioned on 97 baseline-LD (v2.2) model annotations + one TLand organ-specific annotation. We ran this for all trait-organ combinations, resulting in heritability enrichment estimates and standard errors for each combination. An organ was considered prioritized for a given trait if the lower bound of the heritability enrichment estimate’s 95% confidence interval was greater than 0. Specifically, if the contribution of the organ-specific TLand annotation to the trait’s narrow-sense heritability (*h^2^*) is significantly greater than the proportion of SNPs covered by the annotation.

### Chromatin accessibility QTL evaluation and organ prioritization

We downloaded the cluster-specific chromatin accessibility QTL (caQTL) results from Wenz et al. (38) (https://zenodo.org/records/12706263) and obtained coordinates for all tested peaks from personal correspondence with the authors. Cluster 9 was chosen for this evaluation due to its homogenous compositions of biosamples relative to other clusters and a large number of caQTL calls to maximize power for statistical testing. To reduce the number of false positive caQTL calls, we filtered for tested variants within 100bp of their corresponding peak midpoint, hereby referred to as the evaluation set of variants. Positive caQTL calls were defined as variants from this set passing a false discovery rate (FDR) of 0.05. We generated TLand scores and Sei scores for the evaluation sets and calculated the corresponding AUPRC and AUROC metrics with these labels.

Organ prioritization of the cluster-specific caQTLs was done by performing one-sided Welch’s t-tests for each of the 51 organ-specific scores against the caQTL calls. Multiple test correction was applied using the Benjamini–Yekutieli method to control for FDR. The resulting q-values for each organ-specific score were log-transformed and ranked in descending order. The top significant organs represent the most relevant organs for a set of cluster-specific caQTLs.

## Results

### TLand incorporates comprehensive datasets to predict regulatory variants

We developed a new model architecture, TLand, to predict regulatory variants from a comprehensive set of features derived from RegulomeDB (Fig 1b and Supplementary Fig 1). TLand takes input as genomic positions or dbSNP IDs, queries RegulomeDB for features, and then outputs the probability that each variant is a cell-type specific (Supplementary Fig. 1) or organ-specific (Fig. 1b) regulatory variant. Importantly, by combining stacked generalization, feature grouping, and interaction terms in meta-classifier training, the TLand model can incorporate growing training datasets and novel data types into its feature space while combating overfitting, whereas its predecessors cannot.

The continuously growing corpus of genomic data captures an increasingly complete view of human regulatory variation. RegulomeDB recently upgraded to version 2, expanding to >650 million and >1.5 billion genomic intervals in hg19 and GRCh38, respectively (11). The large discrepancy in the data availability between the two assemblies, for example, in ChIP-seq and open chromatin data (RegulomeDB TF ChIP-seq availability in Fig 1a, open chromatin in Supplementary Fig 3, histone ChIP-seq in Supplementary Fig 4), results from better representation of complex variation and correction of sequencing artifacts in the GRCh38 assembly (12). Features used to predict regulatory variants included experimental and computational features derived in GRCh38, the majority of which were derived from

RegulomeDB v2 (Supplementary Table 1). 1,372 DNase-seq BigWig files were quantile-normalized and used to derive five quantile features for each genomic locus. The DeepSEA disease score feature (31) was replaced by its successor Sei (32). The Sei model simultaneously predicted 21,907 binary assay labels, which were dimension-reduced to 40 features representing sequence classes. Assembling deep learning model predictions and training with comprehensive features reduced the variance of our final models and improved generalization to new cell lines and less-studied organs (39,40).

Training data from six common cell lines were concatenated to train an agnostic model, while the specificities of input features determined the cell- or organ-specificity of model predictions. Features were bagged into three biologically meaningful subspaces before each set was fed into an individual base classifier. Subspaces included a generic experimental feature set, a computational feature set, and a cell/organ-specific experimental feature set for regulatory variants.

We adapted the stacked generalization algorithm to stack the output of individual classifiers and use a meta-classifier to compute the final prediction (19). Stacking enables the use of each individual classifier’s strengths by using their outputs as inputs to a final meta-classifier. Cross-validation is required to prevent information leakage during meta-classifier training.

We made several modifications to the algorithm. We used group cross-validation to make base classifiers to learn the regulatory function as the conditional probability between generic features and cell/organ-specific features. Interaction terms were calculated prior to their inclusion in the meta-classifier. We used probability rather than binary classification in the first layer of base classifiers to train the meta-classifier, achieving higher accuracy (41).

### TLand improves regulatory variant predictions

Cell-specific TLand models substantially outperformed state-of-the-art models in predicting unseen cell-line regulatory variants (Fig. 1c and Supplementary Fig. 5). On average, across holdout cell lines, cell-specific TLand models outperformed the previous best model (TURF) with the area under the precision-recall curve (AUPR) and the area under the receiver operating characteristic (AUROC) increasing from 0.389 to 0.471, and 0.729 to 0.776, respectively. While GenoNet outperformed TURF in three held-out cell lines, the observed gains were predominantly attributed to information leakage. This is because the GenoNet models for held-out cell lines, which were initially trained and tested within that held-out cell line, were directly used in the benchmarking. GenoNet performance was used to evaluate the upper limit of the held-out prediction task. Cell-specific TLand models still outperformed GenoNet, improving the average AUPR by 0.09 and the AUROC by 0.04. The high performance of cell-specific TLand can be explained in two ways. First, we derived comprehensive feature sets for predicting regulatory variants from RegulomeDB, categorizing them into experimental feature sets and a complementary non-correlated computational feature set (i.e., deep learning feature set) (Supplementary Fig. 7). The inclusion of experimental features enabled a ∼30% performance gain compared to DeepSEA and Sei models, which use only deep learning features, with average TLand average AUPR=0.471 compared to AUPR=0.336 for DeepSea and AUPR=0.364 for Sei. Second, TLand’s architecture models different biological domains separately and then calculates their conditional probabilities. The meta-classifier finds the best combination of these probabilities and subtracts redundant information through the inclusion of negative coefficients (Supplementary Fig 8 and Supplementary Fig 9) to cancel out noise, thus improving predictions, yielding improvements of 0.10 in average AUPR and 0.04 in average AUROC when compared to TURF (Supplementary Fig 10).

### Organ-specific TLand models address data availability bias

TLand’s flexible architecture enables substituting cell-specific features with organ-specific features, which can then be retrained with organ-specific labels to develop organ-specific TLand models. Organ-specific TLand models can predict organ-specific regulatory variants, even when the organ is represented by only less-studied cell lines or tissues. These models performed better than their cell-specific counterparts in two out of four cell-type specific tasks, specifically in HepG2 (AUPR 0.547 vs. 0.514, AUROC 0.781 vs. 0.692) and MCF-7 (AUPR 0.512 vs. 0.512, AUROC 0.879 vs. 0.840). However, because the GM12878 and K562 cell lines have conflicting labels for the same organ (blood), the blood organ-specific model was evaluated only in the organ-specific (i.e., heart) task, not in either of the cell-line-specific tasks. The superior performance of organ-specific models, even in hold-out cell line tasks, can be attributed to the fact that cell-type specific regulatory variants in HepG2 and MCF-7 are well-represented by corresponding organ-specific regulatory variants within the RegulomeDB database (see Supplementary Fig 11). In the A549 cell line, which is from the least-represented organ (lung), the organ-specific TLand model underperformed the cell-specific TLand model holdout on A549 (AUPR 0.329 vs. 0.347), as the organ model predicted regulatory variants in the lung that were not observed in the A549 cell line. To better evaluate model performance in the lung, the ASB dataset from the second-most representative lung cell line, IMR-90, was added to the holdout dataset. The inclusion of this dataset led to TLand outperforming the cell-specific model when held out on the lung with the IMR-90 dataset (AUPR 0.483 vs. 0.432; AUROC 0.841 vs. 0.814), indicating that the organ-specific TLand model can predict a comprehensive set of organ-specific regulatory variants.

However, adding the second most representative cell line in the embryo, H9, did not improve the TLand organ-specific model when compared to the cell-specific model when holding out on the embryo organ (AUPR 0.536 vs. 0.597). Deep learning features, such as DeepSEA disease impact score (31) and Sei sequence classes (32), were more representative of the cell lines or organs with more data availability. This is because the scores were dimensionally reduced from 919 and 21,907 prediction tasks of experimental assays for DeepSEA and Sei, respectively (including TF and histone ChIP-seq and open chromatin assays). Cell lines and organs with more available data dominated the dimension-reduced results. Thus, we removed deep learning features and developed the organ-specific TLand light model (Supplementary Fig 12) to predict regulatory variants in organs with low data availability, such as the embryo. We defined organs with >=100 available ChIP-seq assays as “high data availability” and those with <100 assays as “low data availability”. We observed that the TLand (full) model consistently outperformed the TLand light model in organs with high data availability (Fig. 1a and c; Supplementary Fig. 5), whereas the light model outperformed the full model in organs with low data availability. For example, the organ-specific TLand light model performed best when holding out the embryo organ (AUPR 0.639, AUROC 0.774). Those findings indicate that TLand and light models are suitable for predicting regulatory variants in organs with low data availability, whereas TLand (full) models are more suitable for organs with high data availability. We then proceeded with our analysis by training the TLand and TLand light models on all available data. We trained an additional TLand model, TLand lightest, in which the organ-specific ChIP-seq features were removed to further reduce bias toward overrepresented organs (benchmarking results shown in Supplementary Table 2).

We also evaluated the TLand models using leave-one-chromosome-out cross-validation (LOCO CV) across aggregated cell line data to ensure that data leakage via Sei features was not the source of TLand’s improved performance (Supplementary Fig 6). The Sei features depend on a 4096-bp DNA sequence as input, with the variant positioned at the center. This allows for train and test set variants to potentially overlap the same input sequence, resulting in data leakage between the train and test sets. This potential data leakage is relevant only to the full TLand models, as the light and lightest models do not use Sei features. We show that the TLand still outperforms other models using LOCO CV, and that organ-specific models substantially outperform the cell-specific model. The complete organ-specific model performs best, consistent with the held-out cell line results, as the blood and liver variants are included in this cell line-aggregated dataset.

Finally, we evaluated our final models with independent MPRA datasets. We observed similar trends, with TLand models outperforming alternative methods (Supplementary Fig. 13). However, MPRA evaluation can only test for a subset of our predicted variants since MPRA variants affect gene expression. In contrast, TLand models predict variants that have any regulatory effects (defined as altering TF binding), which may not affect gene expression.

### TLand prioritizes relevant organs for GWAS traits

To systematically evaluate organ-specific TLand models, we predicted approximately 2 million GWAS SNPs, including those within the same LD blocks, across 51 organs defined by ENCODE (12). We found that TLand predictions were more correlated with TLand light predictions than TLand lightest (e.g., in the heart organ in Supplementary Fig. 14). We manually curated a list of GWAS traits with relevant organs. TLand, TLand light, and TLand lightest models prioritized the relevant organs for 44 GWAS traits (Supplementary Table 3), sourced from the GWAS catalog with an average accuracy of 0.311, 0.340, and 0.466, respectively (Fig. 2b., see the definition of accuracy in Methods). By integrating prioritized organs from two distinct models, TLand and TLand lightest, the average accuracy was increased to 0.482. The overall low accuracy was partially due to uncertainty about whether low scores reflect low data availability or indicate true negatives, with the target organ lacking regulatory function.

**Figure 2.**
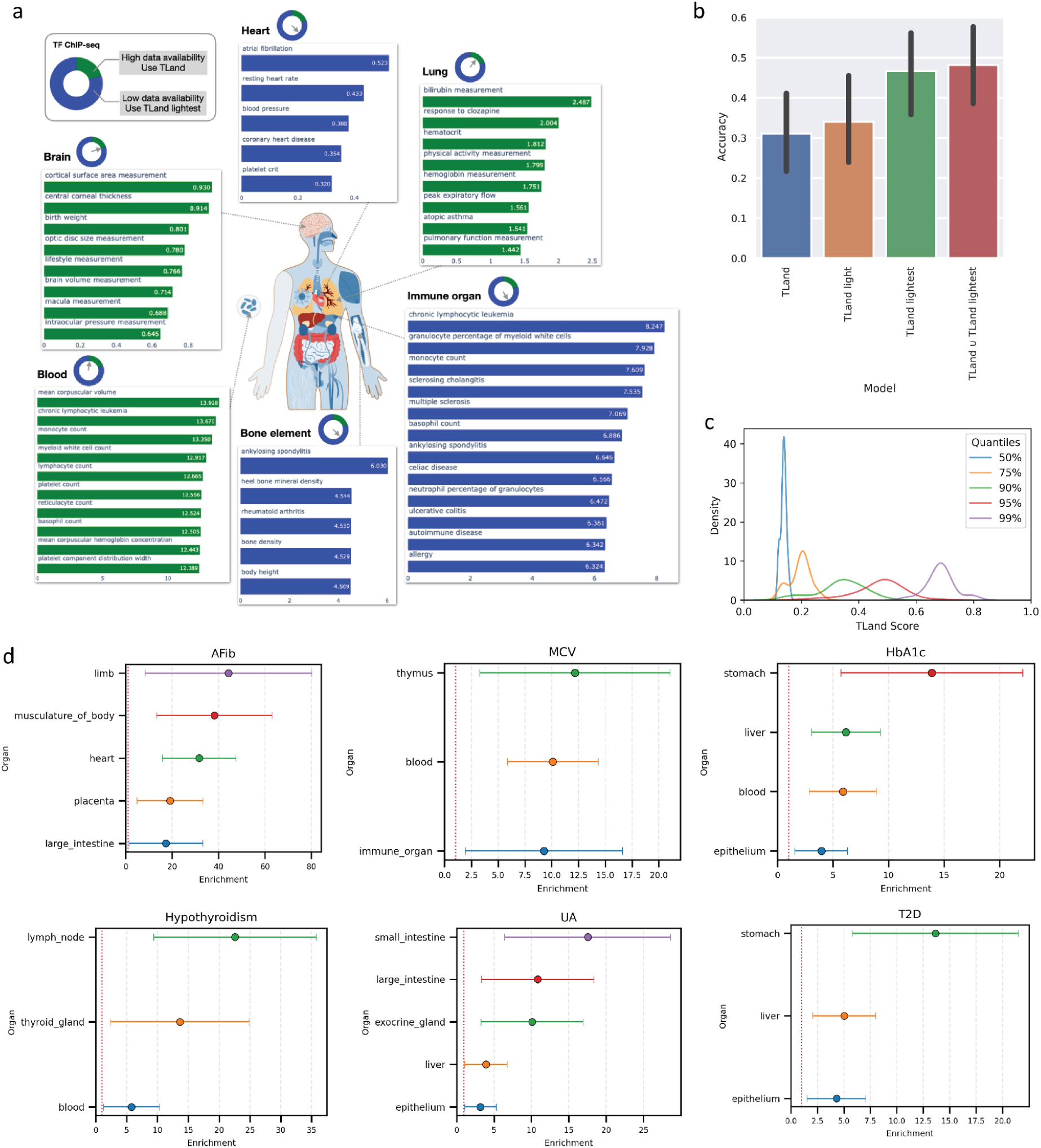
TLand prioritizes relevant organs for GWAS traits. (a) Prioritization of GWAS traits given top-scored variants by TLand and TLand lightest models. TF ChIP-seq data availability plot is re-plotted in the top left where green represents high data availability and TLand was the model used, and blue represents low data availability, and TLand lightest was the model used. Each circle next to the organs indicates the data availability of each organ. The colors of enrichment bars correspond to the model used. (b) Performance of TLand models prioritizing organs. X-axis are TLand models.

To better evaluate TLand models in organs with limited data availability, we focused on the top-scoring variants in those organs and asked whether the top-scoring variants identified by those models were biologically meaningful. We found that top-scoring variants from organ-specific models were enriched for GWAS traits associated with their target organs (Methods, Fig. 2a, and results in Supplementary Table 4). For example, two of the three most-enriched GWAS traits identified by the TLand lightest model were relevant to the heart organ: atrial fibrillation and resting heart (Fig 2a). Originally, TLand lung models were unable to prioritize the lung organ given the corresponding GWAS SNPs for relevant traits (Supplementary Table 3). However, we found that traits such as physical activity measurement and peak expiratory flow were enriched by top-scored variants in the TLand model. Traits that were associated with multiple organs were properly identified by multiple-organ models. For example, both the bone element and the immune TLand lightest models pinpointed ankylosing spondylitis, a type of inflammatory arthritis affecting the spine and large joints (42), as one of the most enriched GWAS traits for their top-scored variants (Fig 2a).

We also explore if organ-specific TLand scores are enriched for heritability in 94 GWAS traits from the UK Biobank. This is a well-powered cohort, allowing us to utilize the stratified LD score regression (S-LDSC) method to estimate the contribution of TLand scores to each trait’s narrow-sense heritability (Methods). We evaluated distributions of TLand scores at the 50, 90, 95, and 99 percentiles across all organs to identify an appropriate score threshold (Fig. 2c). The top 50% of scores are largely between 0.1 and 0.2. The top 5% of scores span a very wide range, from 0.2 to 0.7. The top 1% of scores contains a reasonable distribution of high-scoring variants between 0.6 and 0.8, with the median around 0.7. We thus used a threshold of 0.7 to binarize TLand scores for use in S-LDSC. We computed enrichment scores for each trait-organ combination and identified significantly enriched organs for 71 traits (Methods, Supplementary Table 6 and Supplementary Table 7). Traits were enriched for a median of 3 organs, with stage 4 balding having the highest number of enriched organs at 17. We see that traits largely decompose into their relevant organs. Lymph node and thyroid gland TLand scores are enriched in hypothyroidism, and body musculature and heart TLand scores are enriched in atrial fibrillation (Figure 2d). Liver TLand scores are also expectedly enriched in HbA1c levels and type 2 diabetes (T2D), but stomach TLand scores are too. Gastroparesis - the delayed movement of food from the stomach - is a common complication of T2D (43) and is hypothesized to be due to vagus nerve damage from

T2D-induced hyperglycemia. Still, the cause is ultimately unclear (44). There is also limited research on the genetic factors of gastroparesis. The enrichment of stomach TLand scores in T2D could suggest a pleiotropic model in which T2D-associated variants affect both T2D and stomach traits, explaining the frequent co-occurrence of T2D and gastroparesis. Overall, we show that organ-specific TLand models can accurately prioritize relevant organs for GWAS traits and be used to propose hypotheses for complex genetic risk factors.

Y-axis is accuracy (definition see Methods). (c) Distribution of TLand scores using 50, 75, 90, 95, and 99% quantile thresholds across organs. (d) Significant S-LDSC enrichment results using TLand scores for atrial fibrillation (AFib), mean corpuscular volume (MCV), hemoglobin A1C (HbA1c), hypothyroidism, uric acid (UA), and type 2 diabetes (T2D). Error bars denote 95% confidence intervals.

### TLand partitions caQTLs into relevant organs

We evaluated TLand on chromatin accessibility QTL (caQTL) prediction and ability to recover sample tissue identities from cluster-specific ATAC-seq datasets. Wenz et al. collected 10,293 publicly available ATAC-seq datasets, performed genotyping directly from the ATAC-seq reads, and called caQTLs both globally and on clustered samples (38) (Figure 3a). Clustered samples generally reflected similar tissue types or treatment/disease conditions. We calculated organ-specific TLand scores for these cluster-specific caQTLs and limited the evaluations to tested variants within 100bp of their corresponding peak midpoints to minimize false-positive caQTL calls and ensure that TLand was differentiating between likely causal vs non-causal variants instead of simply predicting presence/absence of peaks at variant positions. TLand blood scores clearly separate non-caQTLs and caQTLs in cluster 9 (Figure 3b), which is largely composed of lymphoblastoid cell lines, and outperform Sei in classifying caQTLs in this cluster (Figures 3c and 3d). For each of the 51 organ-specific TLand scores, we tested if the mean organ-specific score of caQTLs was greater than the mean organ-specific score of non-caQTLs using a one-sided Welch’s t-test. After performing multiple test correction, we ranked the resulting q-values for each of the tests (where each test corresponds to an organ) in order to prioritize organs associated with the set of caQTLs. Because of shared ATAC-seq peaks and variant effects across organs, almost all tests yield highly significant q-values; however, we expect the most relevant organs to be ranked highly. Blood was the top-ranked organ, and the other highly ranked organs (bone marrow, bone element, immune organ, lymph node, thymus) are also largely composed of immune cell types (Figure 3e). We show that TLand can also be used to prioritize variant effects on chromatin accessibility and partition caQTLs into relevant organs, allowing researchers to impute potential cross-tissue effects.

**Figure 3.**
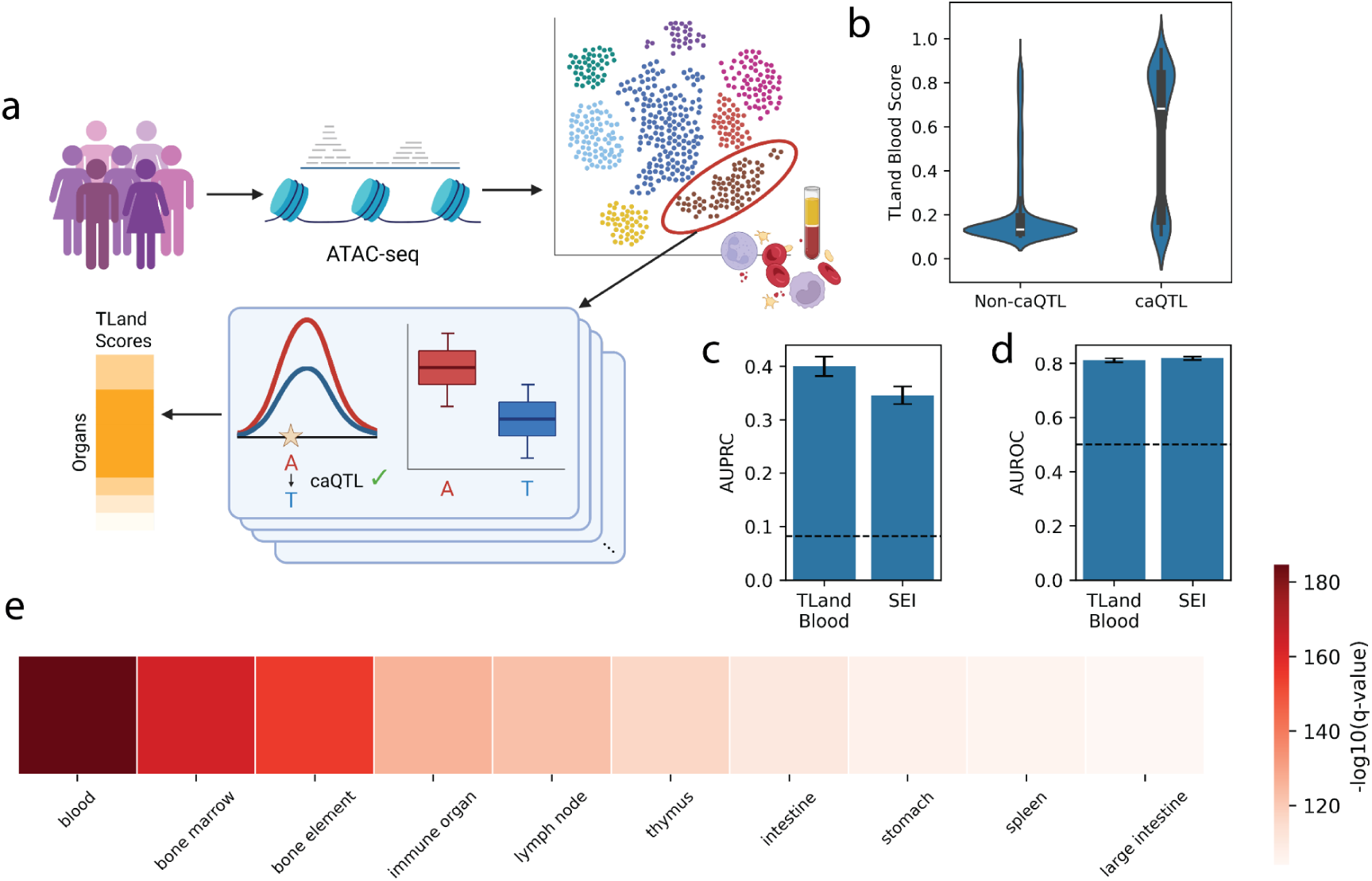
TLand partitions caQTLs into relevant organs. (a) Diagram of organ prioritization of caQTLs. Public ATAC-seq datasets were collected and uniformly processed through a genotyping and caQTL calling pipeline by Wenz et al. They clustered the datasets based on accessibility profiles, and cluster-specific caQTLs were called. We filtered these caQTLs and tested high-confidence caQTLs against high-confidence negatives using organ-specific TLand scores. Multiple test correction was performed, and organs were ranked by q-value. (b) Distribution of TLand blood scores for non-caQTL variants and caQTL variants in cluster 9. (c) AUPRC and (d) AUROC of TLand blood and Sei scores for classifying caQTL variants from (b). (e) Top 10 prioritized organs for cluster 9 caQTLs ranked by p-value.

## Discussion

The identification and characterization of non-coding regulatory variants in the human genome remain a major challenge in genomics. We have developed a novel model architecture, TLand, to accompany the recent advancement of RegulomeDB. TLand uses RegulomeDB-derived features to infer regulatory variants at the cell- and organ-specific levels. By applying stacked generalization to incorporate comprehensive datasets on the GRCh38 reference genome, including experimental and computational features, TLand models consistently outperformed state-of-the-art models in holdout benchmarking. This demonstrates the TLand models’ ability to generalize to new cell lines or organs, a major advancement over the SURF and TURF models they extend. Much of this improvement can be attributed to improved handling of differential data availability. In particular, TLand light and lightest models are applicable to organs with limited data, e.g., the embryo. In contrast, current leading classification models, such as DeepSEA’s disease impact scores, perform poorly in such organs.

One of TLand’s main strengths is that it covers nearly all human tissue types, enabling hypothesizing pleiotropic effects via GWAS-trait heritability enrichment or caQTL organ prioritization. Many existing variant-effect models either capture global, tissue-agnostic pathogenicity or are limited to a few tissue types with maximal data modality coverage. Our generalization approach, based on careful feature bagging and training separate models, enables us to investigate variant effects while accounting for the vast majority of biological contexts.

The TLand models presented here represent a promising foundation for further improvements. These are facilitated mainly by TLand’s ability to ingest new datasets as they are released. We aim to continuously incorporate new datasets into RegulomeDB as they are released, thus extending its performance and applicability as the number and diversity of training datasets grow across different cells, tissues, and organs. Many of these, including gkm-SVM model predictions (45) and Hi-C contact maps (17), are new features for TLand models. TLand’s flexible, modular design allows grouping features into biologically meaningful sets that can be evaluated against one another before making the final decision about whether to include novel features in the model.

One limitation of this study is the limited number of allele-specific binding (ASB) sites in our training dataset. Because of data availability constraints, ASB training datasets were derived from the personal genomes of only six common cell lines. In addition, there is an extensive disagreement among existing ASB prediction methods, further reducing the number of high-confidence training variants. Recently, Adastra, a new database that hosts 652,595 ASBs passing 5% FDR across 647 cell lines and 1,043 TFs, was published (21). However, Adastra predicts ASB using a statistical model, not by using personal genomes directly.

Therefore, it is unclear whether incorporating these ASB predictions in the high-confidence training datasets will yield meaningful improvements to the TLand models. This remains an area of ongoing interest.

## Conclusions

Overall, TLand advances the field of non-coding variant analysis by incorporating comprehensive datasets to predict functional regulatory variants at cell-specific and organ-specific levels. To foster downstream applications, we have provided pre-calculated TLand scores for 30 million genomic variants through the IGVF data portal (See Availability of data and materials section). These scores are being used by the IGVF consortium to prioritize disease and trait-associated variants for functional assays and benchmarking studies. TLand’s flexible models and ability to integrate novel datasets in a principled way significantly improve our ability to identify human regulatory variants genome-wide. This ability will only improve as larger, more comprehensive training datasets spanning additional organs, tissues, cell types, and assays are released.

## List of abbreviations

SNPs: Single-nucleotide polymorphisms
GWAS: Genome-wide association study
ASB: Allele-specific binding
AUROC: Area Under the Receiver Operating Characteristic Curve
AUPR/AUPRC: Area Under the Precision Recall Curve
MPRA: Massively parallel reporter assay
FDR: False discovery rate
LD: Linkage disequilibrium
MHC: Major histocompatibility complex
caQTL: Chromatin accessibility quantitative trait locus
TF: Transcription factor
LOCO CV: Leave-one-chromsome-out cross-validation
S-LDSC: Stratified linkage disequilibrium score
T2D: Type 2 diabetes
AFib: Atrial fibrillation
MCV: Mean corpuscular volume
HbA1c: Hemoglobin A1c
UA: Uric acid

## Declarations

### Ethics approval and consent to participate

Not applicable

### Consent for publication

Not applicable

### Availability of data and materials

Scripts to reproduce the analyses can be found at https://github.com/rnsherpa/tland_supplementary. A Snakemake workflow to run TLand predictions on input variants is available at https://github.com/rnsherpa/TLand-predict. The datasets supporting the conclusions of this article are available on Zenodo at https://doi.org/10.5281/zenodo.18341395. TLand predictions for 30 million variants are available on the IGVF data portal at https://data.igvf.org/prediction-sets/IGVFDS0038SURQ/. caQTL data from Wenz et al. are available on Zenodo at https://doi.org/10.5281/zenodo.12706262.

### Authors’ contributions

NZ and SD developed the model. RNS developed the Snakemake workflow. NZ, RNS, and KBH performed the model evaluations and downstream analyses. NZ, RNS, and APB contributed to interpretation of the results. NZ, RNS, and APB wrote the manuscript with input from all authors. APB supervised the work.

## Acknowledgements

We thank Christopher Castro, Benjamin Hitz, Shengcheng Dong, and Zhewei Shen for their assistance in data submission to the IGVF data portal.

