## Supplementary Materials for "Organ-specific prioritization and annotation of non-coding regulatory variants in the human genome"

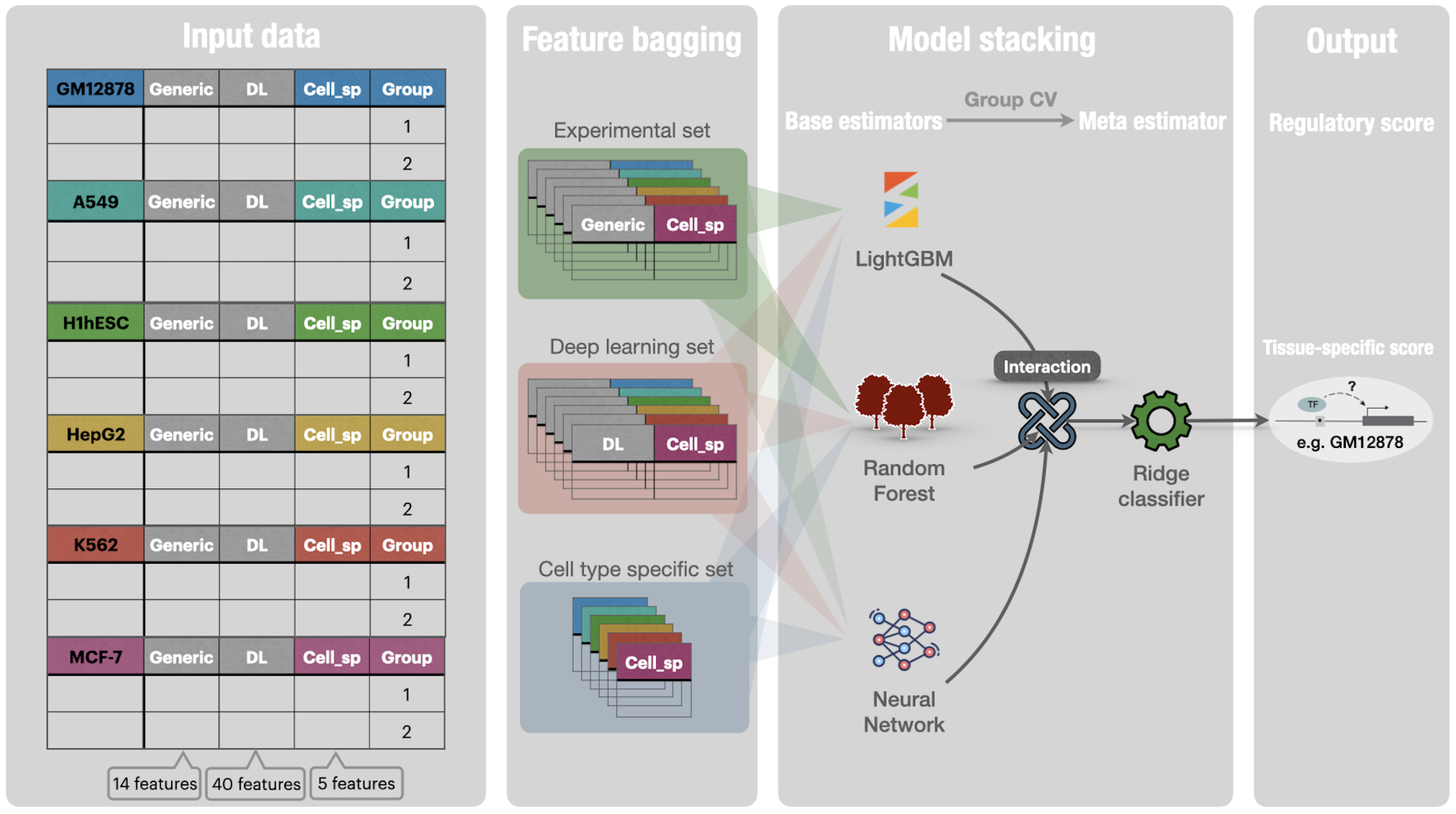

**Supplementary Figure 1. Cell-specific TLand architecture.** Cell-specific TLand was trained to predict human regulatory variants in a cell-specific manner by using RegulomeDB-derived features.

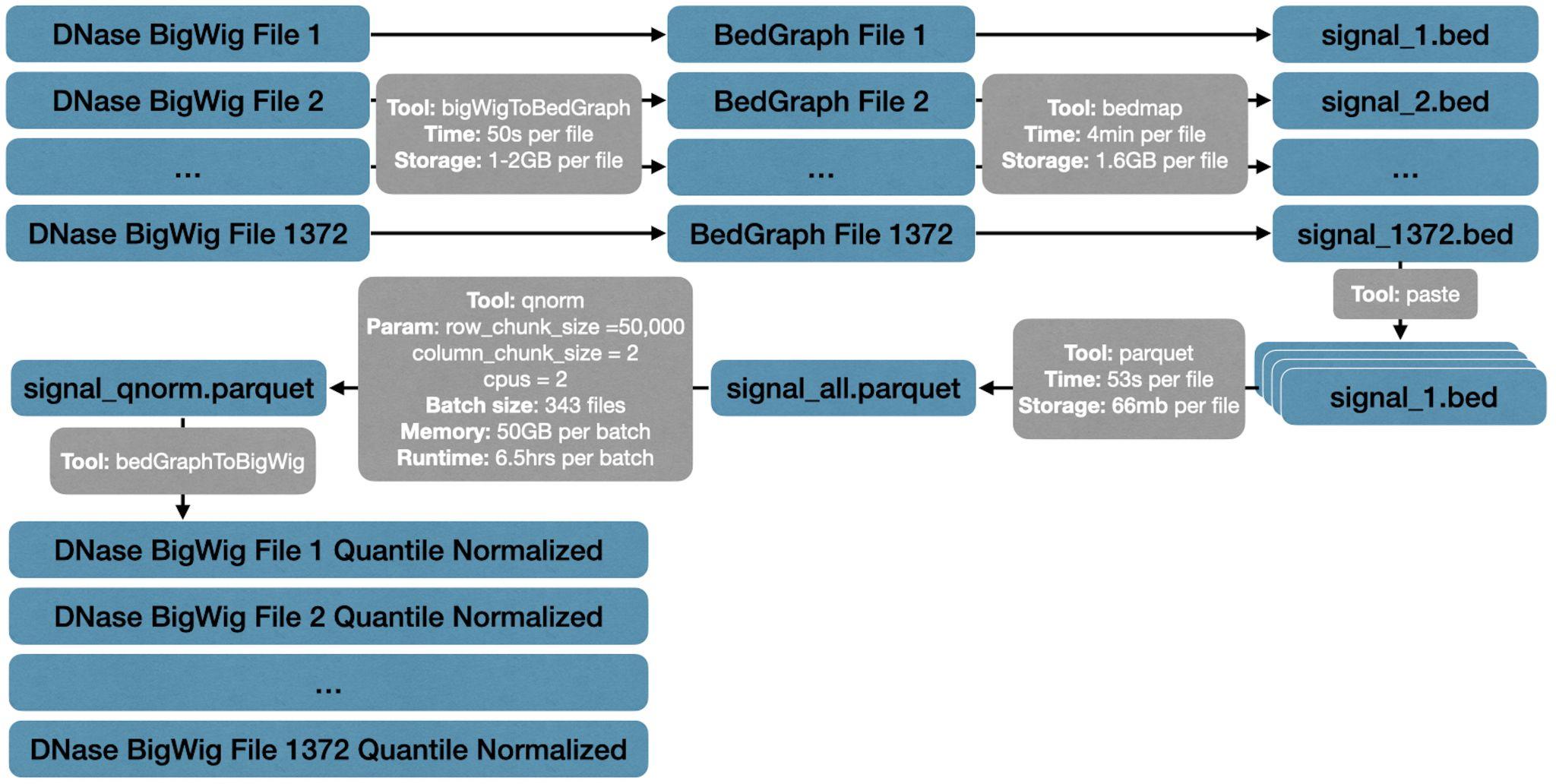

**Supplementary Figure 2.** **Quantile normalization pipeline.** We designed a pipeline to quantile normalize 1372 DNase BigWig files in a bin size of 10 base pairs. Important details to reproduce, including tools, parameters of tools, time, and storage it took, are all specified in the figure.

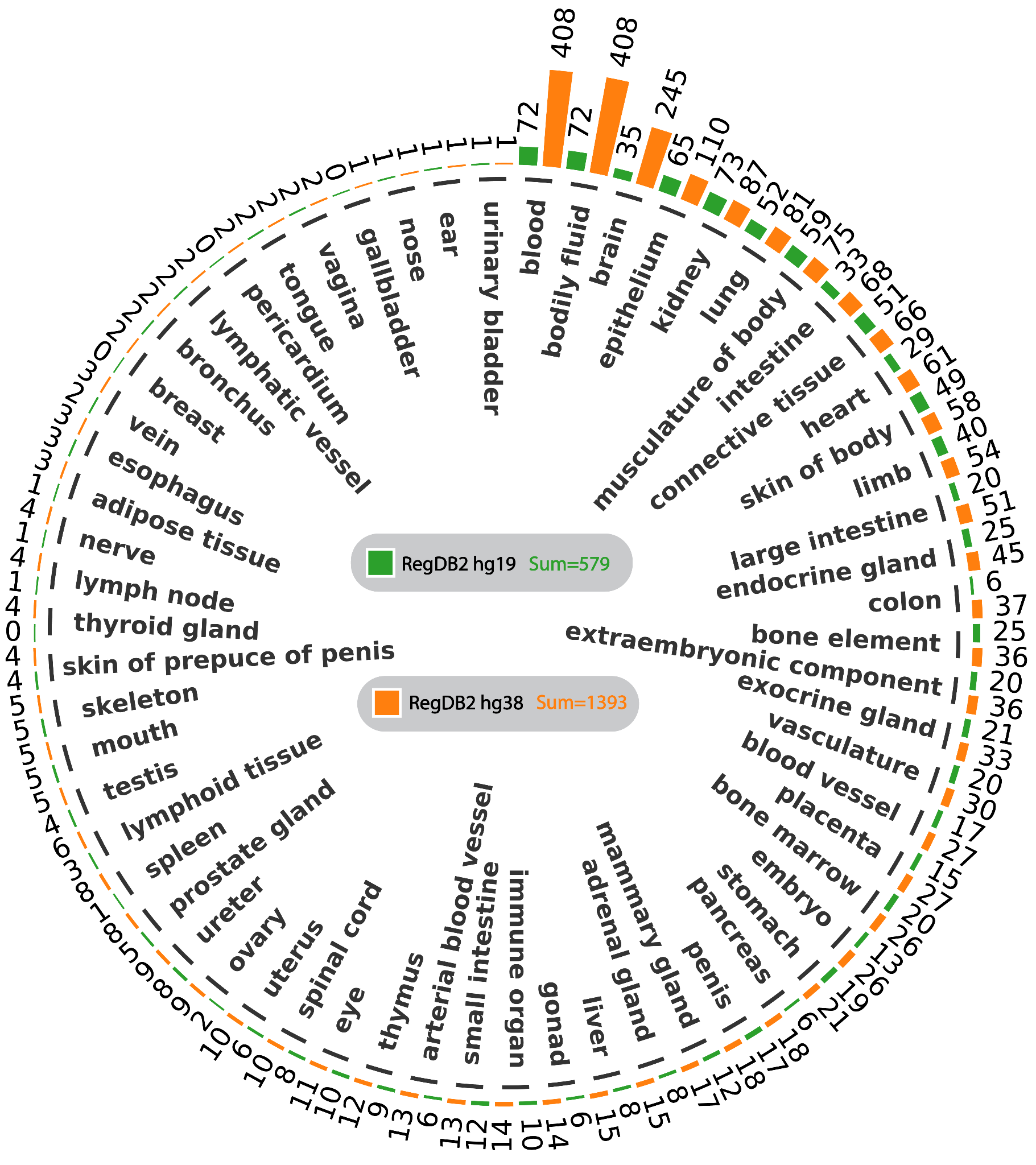

**Supplementary Figure 3. DNase-seq data availability across organs on RegulomeDB v2.** Green bar plots represent counts on GRCh38. Orange bar plots represent counts on hg19. The total number of counts for each assembly is in the middle of the gray box. Notice that the summation is not simply adding all numbers together due to some cell lines having multiple corresponding organs.

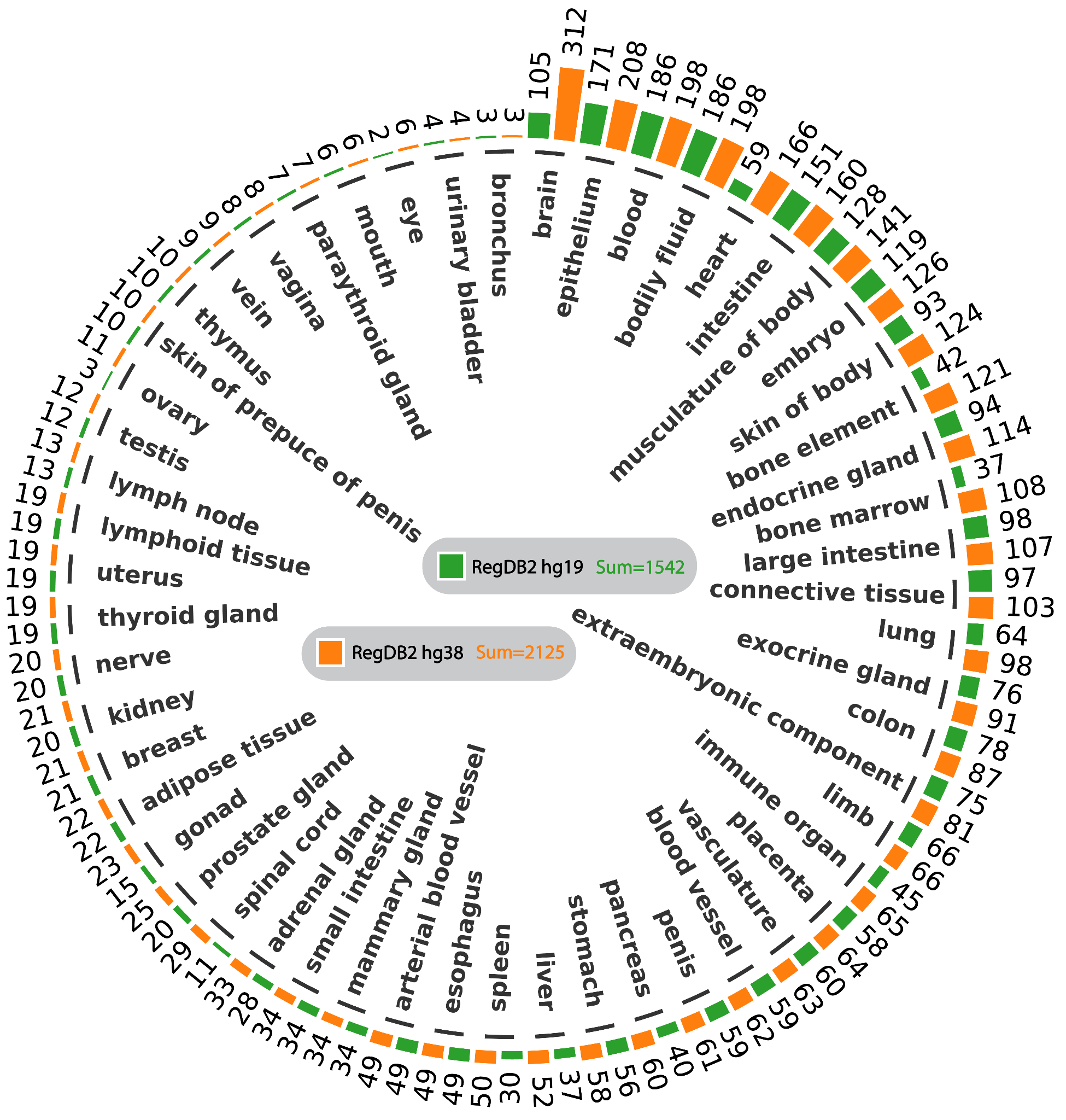

**Supplementary Figure 4. Histone ChIP-seq data availability across organs on RegulomeDB v2.** Histone ChIP-seq data includes five histone marks: H3K4me1, H3K27ac, H3K36me3, H3K4me3, and H3K27me3. Green bar plots represent counts on GRCh38. Orange bar plots represent counts on hg19. The total number of counts for each assembly is in the middle of the gray box. Notice that the summation is not simply adding all numbers together due to some cell lines having multiple corresponding organs.

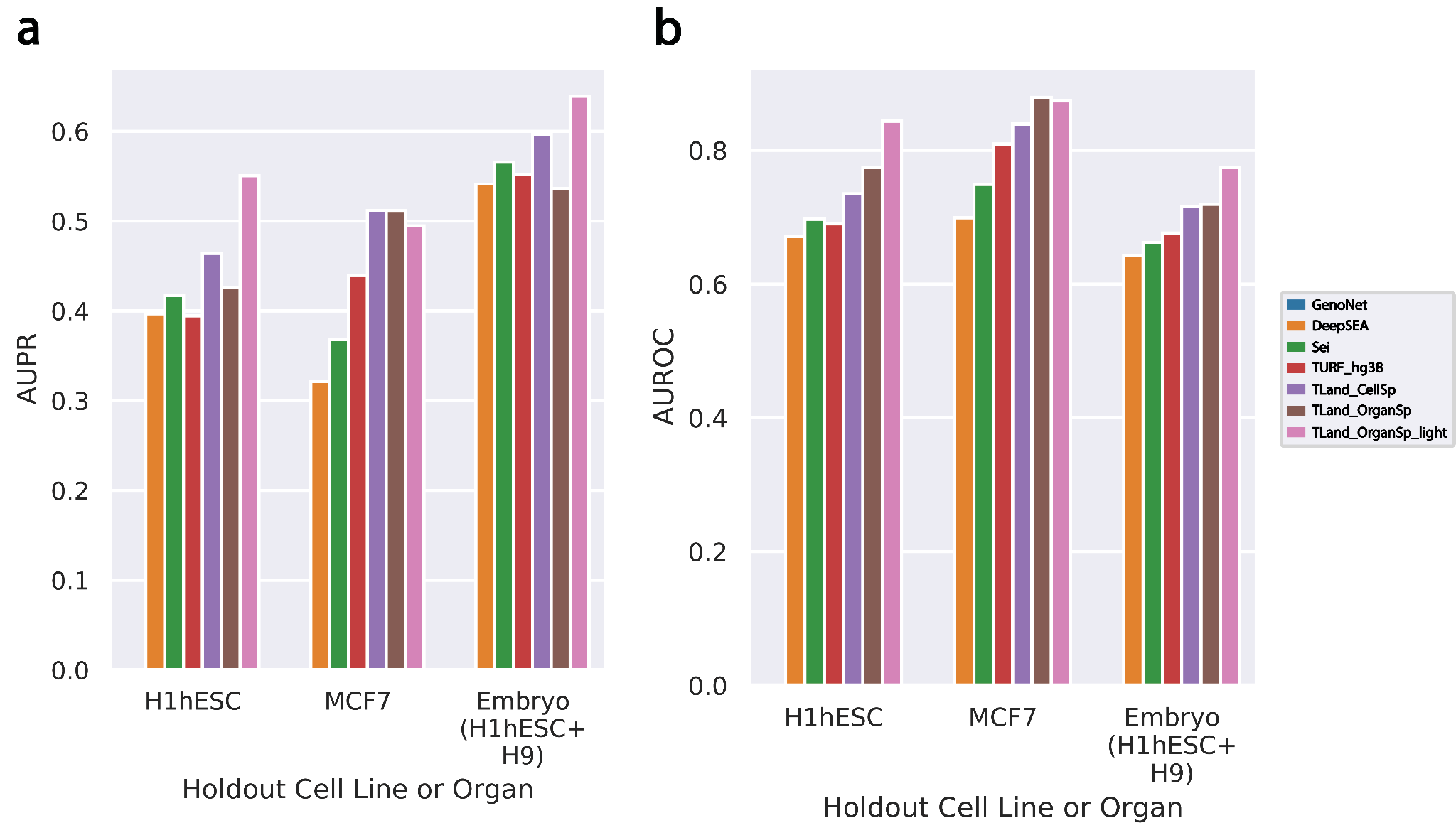

**Supplementary Figure 5. Benchmarking models on H1, MCF7, and embryo.** X-axis is holdout cell lines or organs. Y-axis is AUPR on the left panel and AUROC on the right panel.

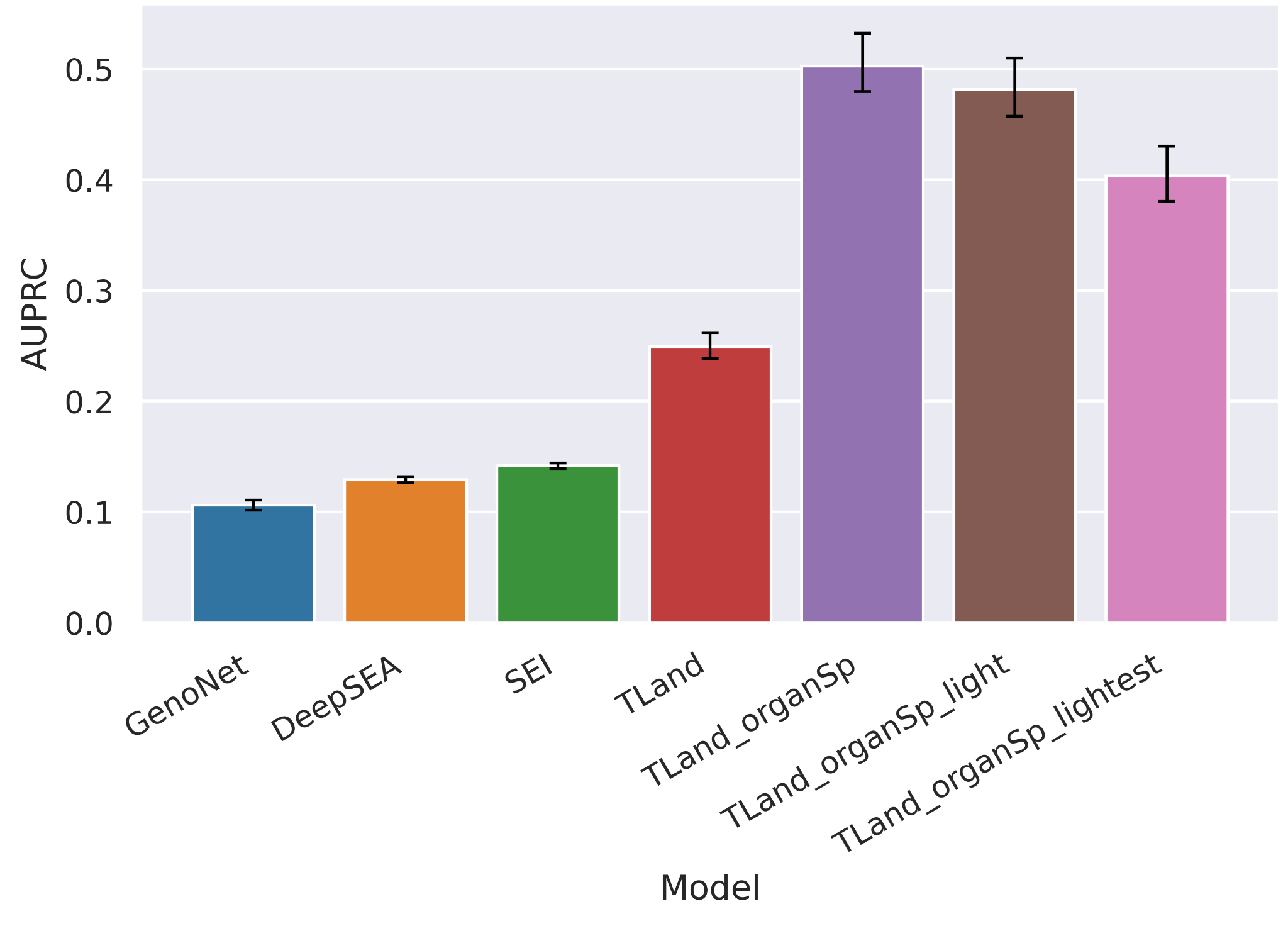

**Supplementary Figure 6. Benchmarking models on aggregated cell line data using leave-one-chromosome-out cross-validation.** X-axis is models. Y-axis is AUPRC. Error bars denote bootstrapped 95% confidence intervals.

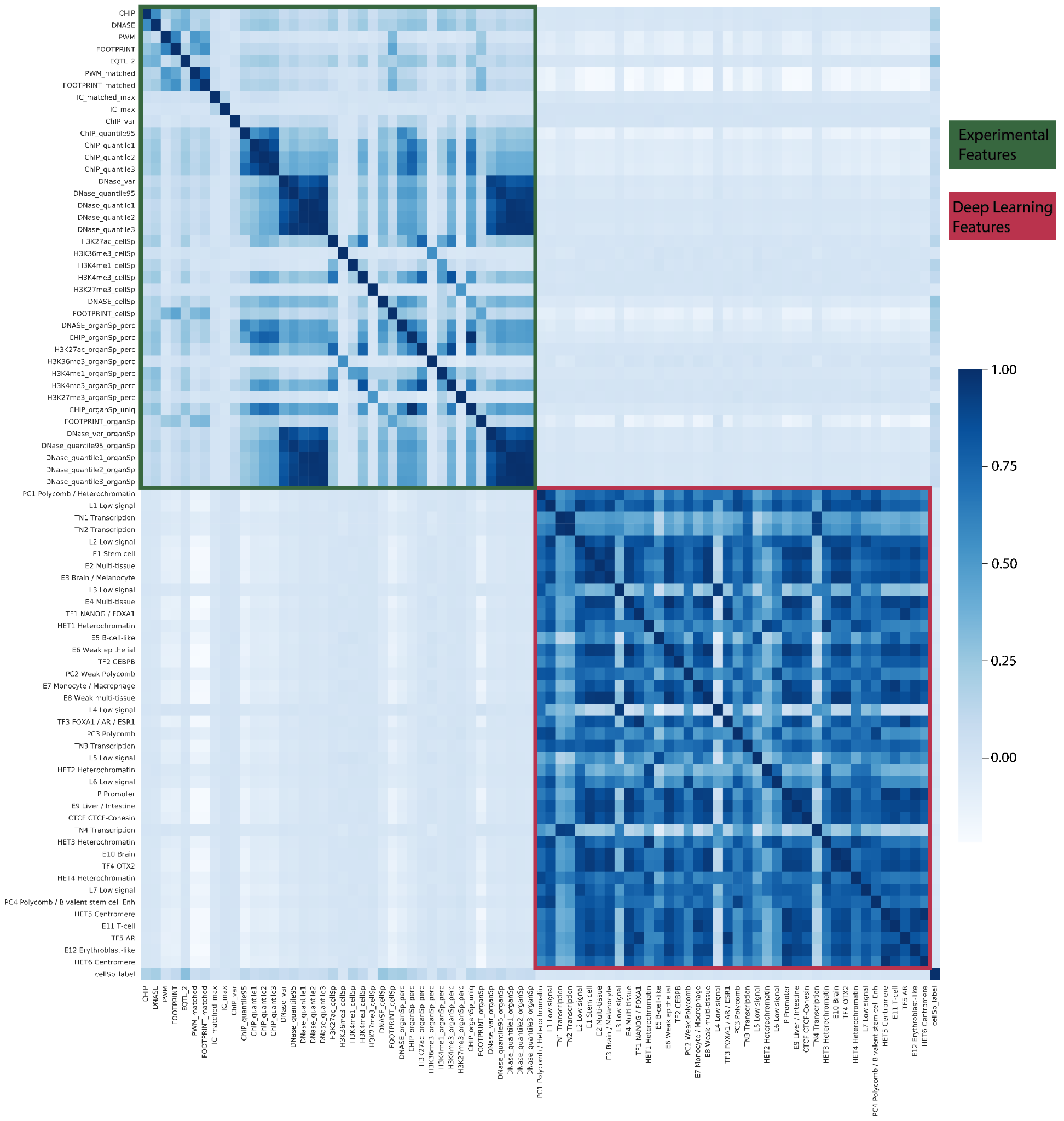

**Supplementary Figure 7. Correlation between deep learning features and experimental features in GM12878 cell line.** Both axes are features derived from RegulomeDB for the GM12878 ASB dataset. The heatmap was calculated as the correlation between those features, ranging from [-1, 1]. The green box represents the features belonging to the experimental feature set. The red box represents the features belonging to the deep learning feature set.

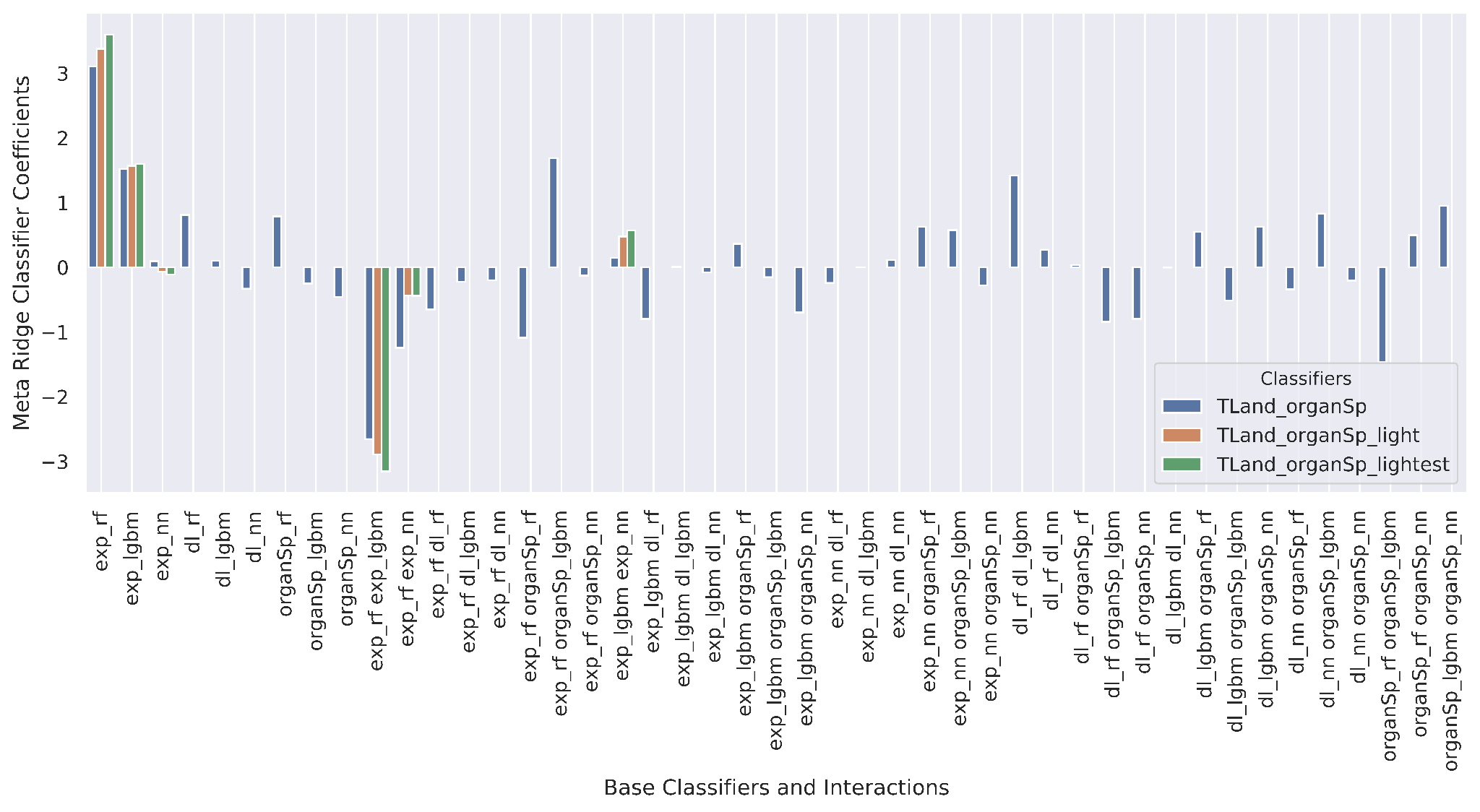
**Supplementary Figure 8. Ridge classifier coefficients.** The X-axis is base classifiers and their interaction term from the final TLand models trained on all data. Y-axis is the coefficient of the meta-ridge classifier. Blue represents TLand organ-specific model. Orange represents TLand organ-specific light model which did not have deep learning features (i.e. no *dl* base classifiers nor *organSp* base classifiers). Green represents TLand organ-specific lightest model which removed organ-specific ChIP-seq features from the light model.

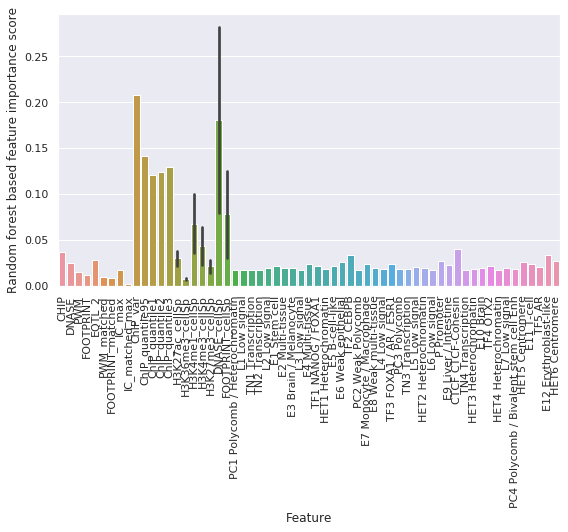

**Supplementary Figure 9. Random forest feature importance scores.** X-axis is a feature in one of the random forest models from the final TLand models trained on all data. Y-axis is feature importance score. The average was taken for cell-specific features since they are in both the experimental feature set and the cell-specific feature set.

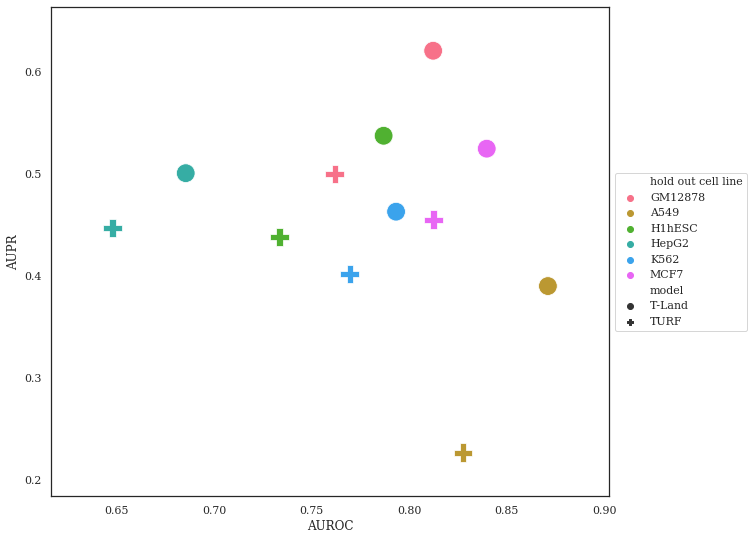

**Supplementary Figure 10. Benchmarking TLand and TURF on hg19.** X-axis is AUROC, and Y-axis is AUPR. Colors represent holdout cell lines. Shapes represent models. Specifically, the circle represents the cell-specific TLand model, and the plus sign represents the TURF model.

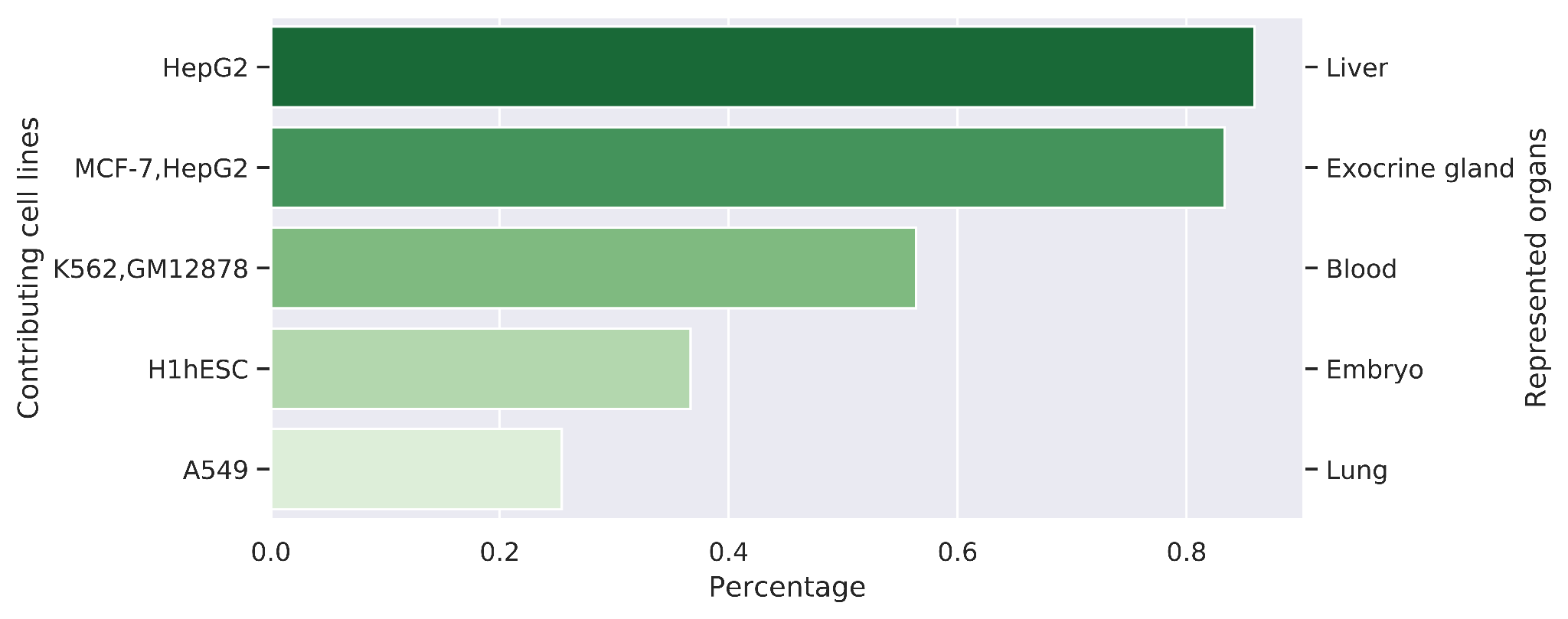

**Supplementary Figure 11. Organ representability by cell lines on RegulomeDB v2.** X-axis is the representability percentage of organs, which indicates how well each organ is represented in the RegulomeDB by the contributing cell lines. The higher the representability, the darker the green color. Y-axis on the left is contributing cell lines. Y-axis on the right is the represented organs.

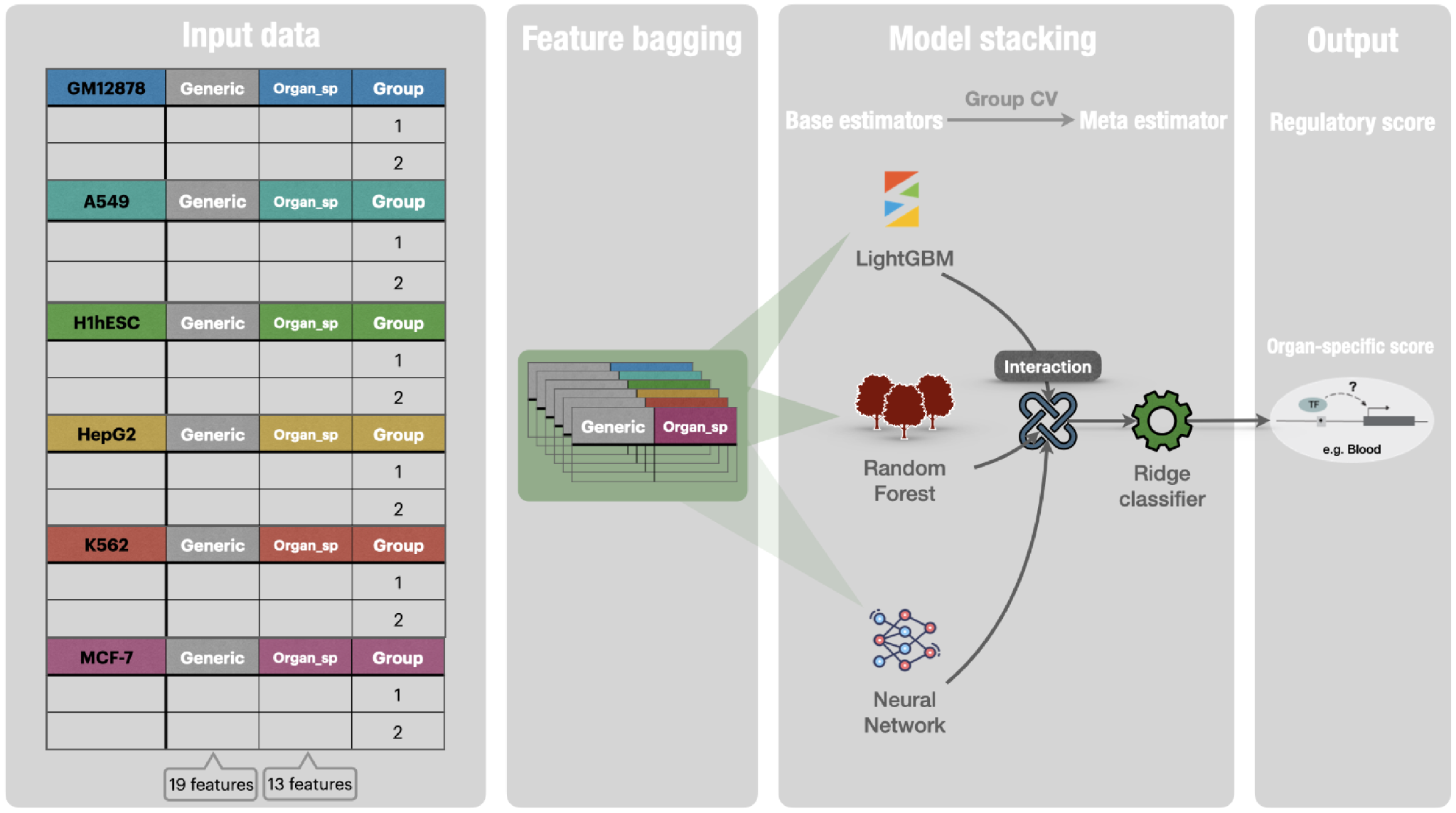

**Supplementary Figure 12. Organ-specific TLand light architecture.** Organ-specific TLand light was trained to predict human regulatory variants in an organ-specific manner by only using RegulomeDB-derived experimental features.

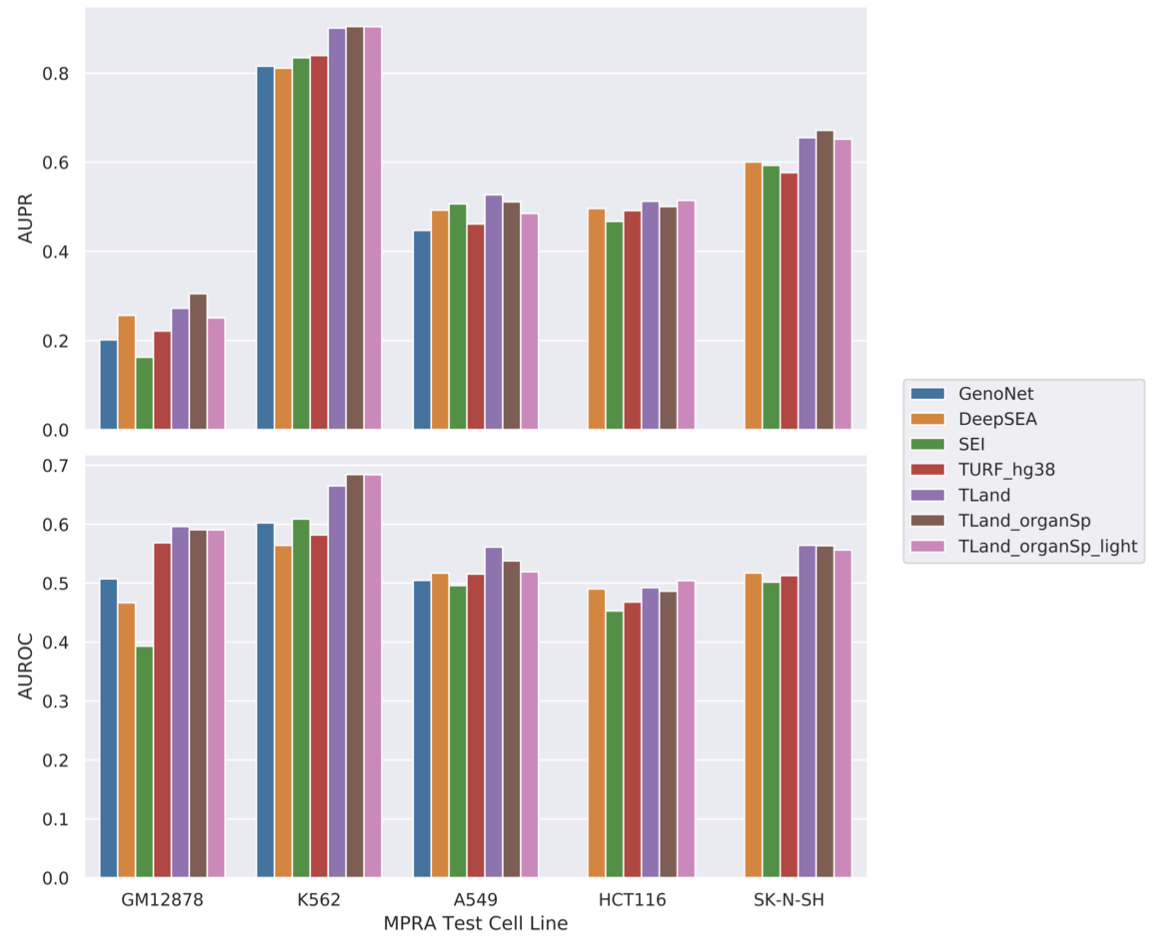

**Supplementary Figure 13.** Benchmarking TLand performance by AUROC and AUPR with MPRA datasets. X-axis is MPRA test cell lines. Y-axis is AUPR on the top panel and AUROC on the bottom panel.

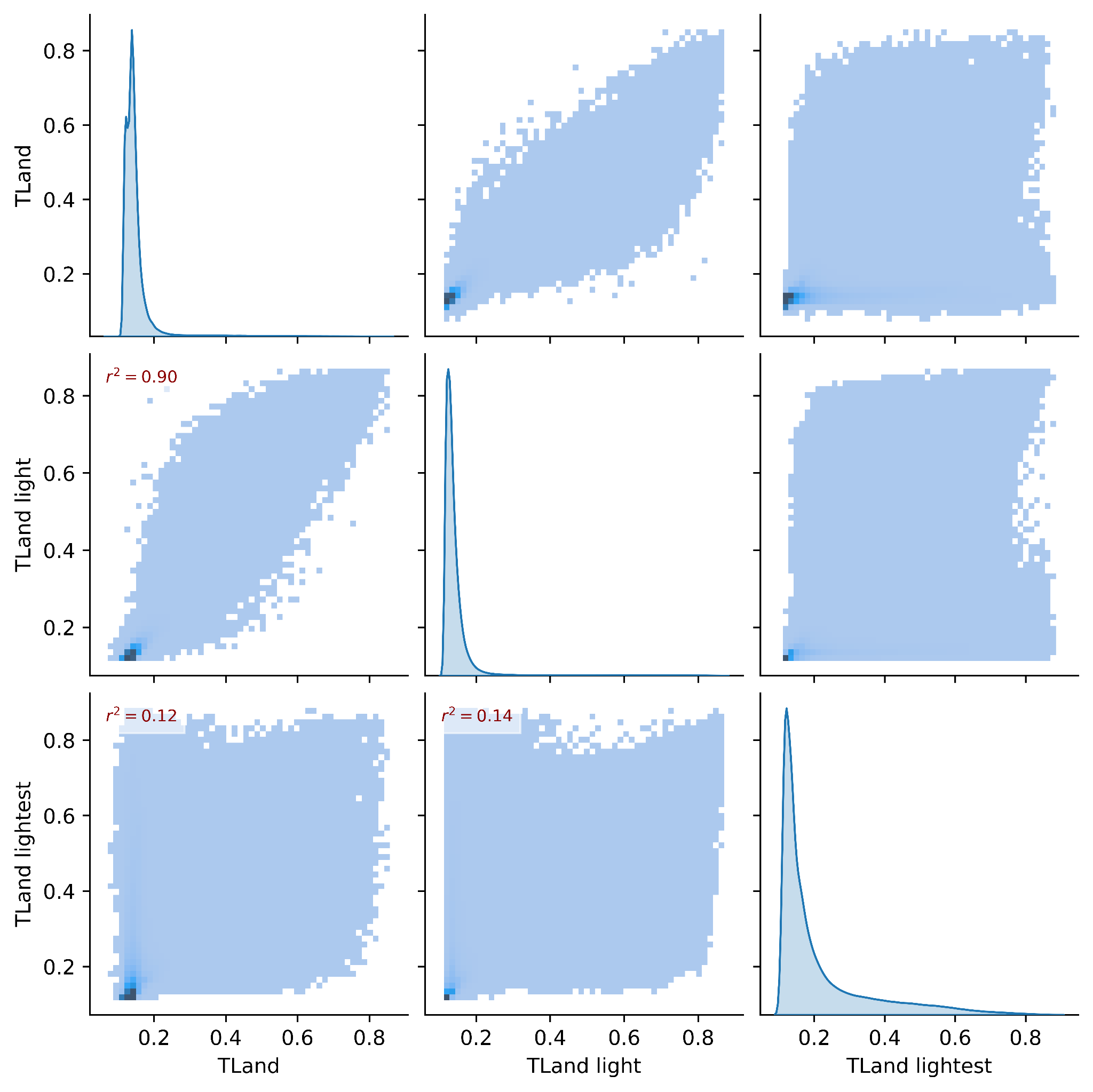

**Supplementary Figure 14. Pairplot of TLand model prediction scores in the heart.** X-axis and Y-axis are TLand, TLand light, and TLand lightest model prediction scores in order.
